# SUGP1 associates Y-box protein to regulate piRNA biogenesis in Bombyx mori

**DOI:** 10.1101/2025.10.09.681363

**Authors:** Dan Liu, Yongkang Guo, Mengting Shen, Jing Lv, Peng Wei, Ling Jia, Sanyuan Ma

**Author notes:** These authors contributed equally to this work (Dan Liu, Yongkang Guo).

## Abstract

PIWI-interacting RNAs (piRNAs) are critical for transposon silencing and genome integrity, as well as gene expression regulation and antiviral immunity in metazoans, yet the molecular mechanisms governing their biogenesis remain incompletely understood. The participation of the splicing-associated process in piRNA biogenesis has been emphasized in multiple species, but the key factors and mechanisms remain elusive. Here, we identified SUGP1 (SURP and G-Patch Domain Containing 1) in BmE (a unique model cell system with a complete piRNA biogenesis pathway) as a key splicing factor that functions in piRNA biogenesis. Through CRISPR-Cas9-mediated gene knockdown in cultured cells combined with RNA-seq, small RNA-seq, and IP-mass spectrometry (IP-MS), our results reveal that SUGP1 deficiency disrupts piRNA accumulation, alters mature piRNA length distributions, and activates transposon expression. Immunofluorescence and Western blot (WB) analyses further demonstrate that SUGP1 interacts with Y-box protein (YBP), which is key regulators of RNA metabolism. Functional validation in Drosophila SUGP1-RNAi lines highlights evolutionary conserved and species-specific roles of SUGP1 in piRNA maturation. Collectively, our data uncover a dual role for silkworm SUGP1 in coordinating YBP-dependent piRNA biogenesis, thus elucidating a novel mechanistic framework for piRNA pathway regulation. Our work also underscores the silkworm as a unique model for studying non-canonical piRNA biogenesis mechanisms, with implications for treating transposon dysregulation-linked diseases.

**Author summary:** Disruption of piRNA synthesis leads to abnormal consequences such as transposon de-repression, posing a significant threat to genomic stability. It is therefore essential to in-depth analysis of the piRNA biosynthetic mechanism. Previous studies have highlighted the involvement of splicing-related processes in piRNA biosynthesis, yet key factors and mechanisms remain poorly understood. This study employed the silkworm cell system, which possesses a complete piRNA biosynthetic pathway. Utilizing CRISPR-Cas9-mediated gene knockout technology combined with RNA sequencing, small RNA sequencing, and immunoprecipitation-mass spectrometry (IP-MS), we discovered that SUGP1 deficiency disrupts piRNA accumulation, alters the length distribution of mature piRNAs, and activates transposon expression. Furthermore, immunofluorescence and Western blot analyses confirmed an interaction between SUGP1 and Y-box proteins (YBPs), key regulators of RNA metabolism. Functional validation using Drosophila SUGP1-RNAi lines revealed that SUGP1 plays dual roles in piRNA maturation. Our study reveals that the silkworm scissor-related factor SUGP1 has dual functions in coordinating YBP-dependent piRNA biosynthesis.

## Introduction

Transposable elements (TEs) constitute a major threat to genomic integrity, as they account for a significant proportion of eukaryotic genomes (10% in fish, 45% in humans, and ¿80% in plants) [1, 2]. Effectient suppression of transposons activity is enssential to maintain genome stability and normal physiological functions. piRNAs, an key component of non-coding RNA, were first identified in germ cells and proved to interact with PIWI proteins to inhibit transposon activity [3]. Furthermore, recent studies have highlighted additional roles for piRNA in processes such as gene expression regulation, translation maintenance, antiviral defense, and sex determination [4–11].

Given the multifunctional roels of piRNAs, extensive progress has been made in elucidating their biogenesis mechanisms. PiRNAs are primarily transcribed from genomic region called piRNA clusters, which are classified into unidirectional and bidirectional clusters. Transcripts from unidirectional clusters undergo conventional mRNA-like processing, including promoter-driven transcription, capping, polyadenylation, and splicing. In contrast, bidirectional clusters initiate transcription via heterochromatin-mediated recruitment of transcription factors and lack polyadenylation and splicing steps [12–15]. Following transcription, precursor RNAs exit the nucleus via nuclear pore complexes, aided by the THO complex and other cofactors, and undergo primary processing in the cytoplasm to mature piRNAs [16].

In somatic cells, precursor transcripts are processed in perinuclear compartments such as Yb bodies and P bodies. Key enzymes like Zuc mediate 5’ end cleavage to create a monophosphate moiety, while Hen1 ensures 2’-O-methylation. Mature piRNAs are then loaded onto PIWI proteins to execute their functions [17–21]. In germ cells, piRNA precursors undergo the ping-pong amplification cycle in the nuage, a process requiring two PIWI proteins (e.g., Aub/Ago3 in Drosophila and Siwi/BmAgo3 in silkworm). Antisense-strand-derived piRNAs bound to Aub/Siwi recognize and cleave transposon mRNAs, generating sense-strand precursors that load onto Ago3/BmAgo3. The resulting piRISC complexes then cleave complementary antisense transcripts, creating new antisense piRNAs that rejoin the cycle, enabling exponential amplification [22–24]. This cycle depends on auxiliary factors such as Vasa/DDX43 for cleavage product release [25, 26], Trimmer for 5’ cleavage specificity [27], and Spn-E/Qin for cycle regulation [28, 29]. Additionally, piRNA-PIWI complexes may transport to mitochondria and P-bodies via Gasz, Mael, and other factors for further maturation [30–33].

Despite extensive progress in elucidating piRNA biogenesis mechanisms, significant gaps persist in understanding the contributions of RNA processing pathways to this process. Genome-wide screening results for piRNA biosynthesis-related genes across species such as Drosophila and Caenorhabditis elegans revealed an consistent enrichment of splicing-related genes [34–36]. However, key questions remain unresolved: whether splicing-related genes directly participate in the piRNA biogenesis, and how splicing machinery mechanistically influence piRNA production. To address thses gaps, our study focused on silkworm SUGP1 (SURP and G-Patch Domain-Containing Protein 1), a critical regulator of RNA processing and stability. SUGP1 facilitates proper RNA splicing via its SURP and G-patch domains, ensuring functional mRNA generation post-transcription [37] . This protein’s biological importance is further underscored by its interaction with SF3B1, a key splicing factor implicated in cancer progression [38–40] . Thus, SUGP1 represents a promising candidate to bridge the gap between splicing mechanisms and piRNA biogenesis.

The silkworm emerges as an ideal model cultured cell system for studying piRNA biogenesis due to its compact genome coupled with a complete piRNA biosynthetic pathway and rich piRNA repertoire [41, 42]. To dissect SUGP1’s role in this process, we employed a multifaceted experimental strategy using CRISPR-Cas9-modified silkworm cells. Specifically, 1) RNA-seq and small RNA-seq analyses revealed SUGP1-dependent alternations in mature piRNA length, sequence diversity, abundance, and splicing site utilization; 2) IP-MS, western blotting (WB), and immunofluorescence experiments identified SUGP1’s physical interaction with Y-box protein, suggesting a functional link to piRNA processing; and 3) comparative studies using Drosophila SUGP1-RNAi lines demonstrated species-specific differences in piRNA biogenesis pathways. Together, our findings establish a novel regulatory axis for SUGP1 in modulating piRNA biogenesis, expanding our mechanistic understanding of this process and highlighting the evolutionary diversity of splicing-related factors in piRNA pathways.

## Results

### 0.1 Functional disruption of SUGP1 leads to transposon and viral derepression

We start the investigation of the function of SUGP1 with knocking out experiment using CRISPR-Cas9 technology in BmE cells (S1 FigH). Following generating the SUGP1 knocked-out cell line, which was confirmed by target sequencing, we observed impaired splicing of target genes and significantly reduced cellular viability compared to controls, supporting its evolutionary conserved canonical role in mRNA splicing processes (S1 FigA-G). Next, to explore whether SUGP1 is involved in piRNA biogenesis, we subjected the SUGP1-KD cells to RNA-seq. Transcriptomic analysis suggests a pivotal regulatory role of SUGP1 in piRNA biosynthesis from three different perspectives: 1) Corroborated by qPCR validation the SUGP1-KO cells harbor sharply increased Bombyx mori late virus (BmLV) and nuclear polyhedrosis virus (BmPnV) loads(Fig 1A-B; S1 FigI-K), in agreement with that inhibition of piRNA/siRNA pathways is known to derepress endogenous viruses that remain latent under normal conditions(Fig 1C-E) [43]; 2) SUGP1-KO significantly derepressed a large number of transposons (Fig 1F); and 3) The mRNA levels of Masc, which is strictly silenced by its antagonistic piRNA (Fem) through ping-pong cycle [44, 45], were significantly upregulated (Fig 1G), consistent with piRNA pathway dysfunction. Despite parallels to Drosophila studies, where splicing factor RNPS1 represses piRNA biogenesis by modulating Piwi expression [46], SUGP1-KD did not alter mRNA levels of silkworm piRNA pathway components Siwi or BmAgo3(Fig 1H-I). Collectively, these findings confirm SUGP1’s dual roles in mRNA splicing and piRNA biosynthesis. The distinct mechanistic divergence between Drosophila and silkworm highlights evolutionary flexibility in splicing-piRNA crosstalk, warranting further exploration of shared versus species-specific regulatory nodes.

**Fig 1.**
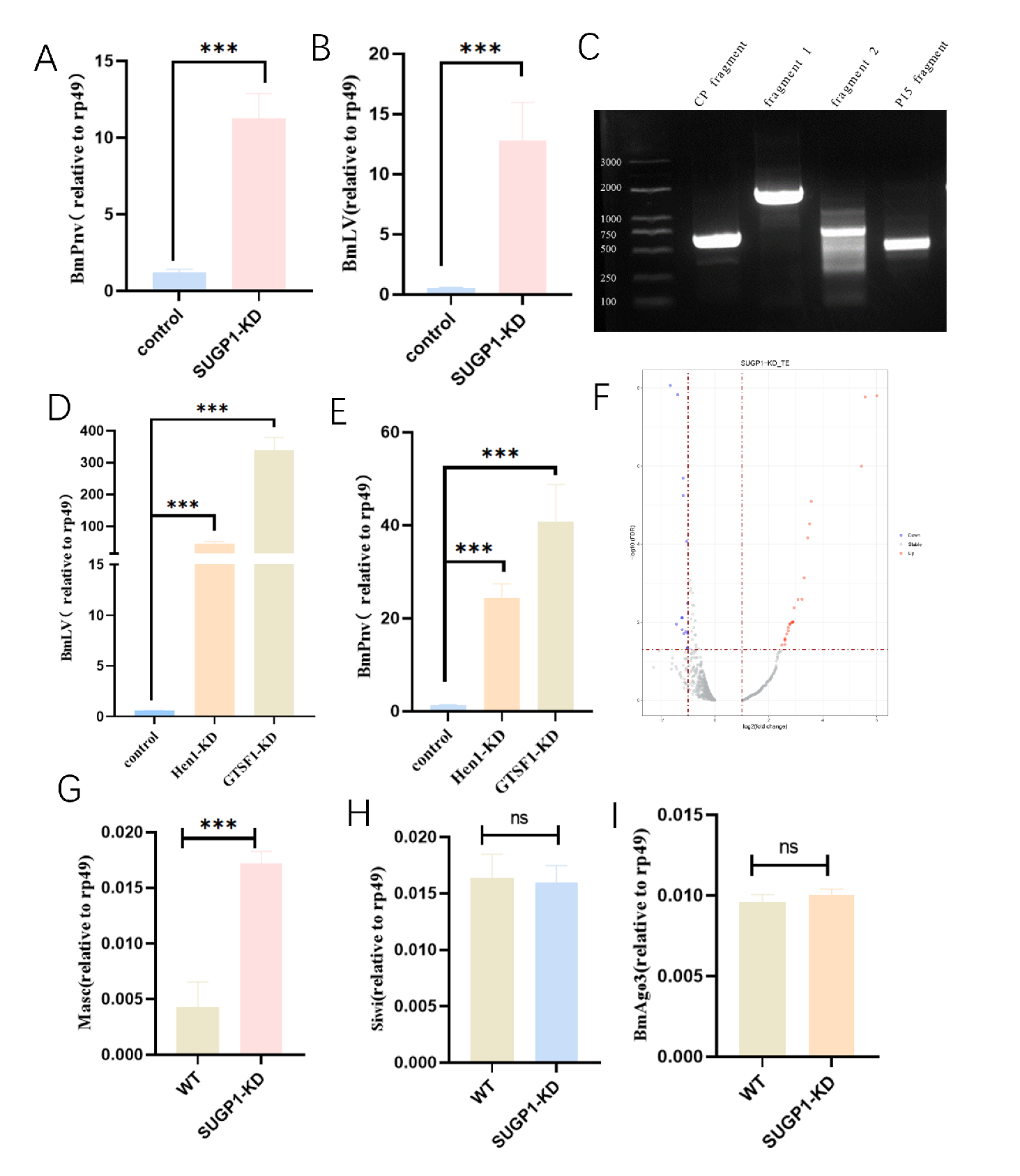
Functional disruption of SUGP1 leads to transposon and vira derepression. (A-B)Changes in BmLV/ BmPnV viral load after knockdown of SUGP1. (C)Amplification of BmLV viral gene fragments in BmE cells of the silkworm. (D-E)Changes in BmLV/ BmPnV viral load after knockdown of core components of the piRNA biosynthetic pathway. (F)Changes in transposon expression after SUGP1 knockdown. (G-I)Changes in Masc/Siwi/BmAgo3 expression after SUGP1 knockdown.

### 0.2 SUGP1 impacts piRNA biogenesis in silkworms through altered RNA processing

To elucidate the role of SUGP1 in piRNA biogenesis, we performed small RNA-seq on SUGP1-KO cells. The length distribution analysis revealed a significant reduction in small RNA populations, particularly with pronounced downregulation of 28 nt and moderate suppression of 32 nt species (Fig 2A). Given that mature silkworm piRNAs are predominantly 27-28 nt, we hypothesized that SUGP1 depletion disrupts the formation of normally sized piRNAs. To dissect the mechanisms driving this length shift, we utilized the PiRBase database to extract all piRNA candidates from the small RNA-seq data. Using the piRNAs identified in the PiRBase database as a control, we aligned the 3–26 nt regions of piRNAs from the sequencing data to the control piRNAs to further analyze the displacement of piRNAs (Fig 2B). Comparative analysis demonstrated that 28 nt piRNA counts and diversity were sharply reduced post-SUGP1-KD, whereas 32-34 nt piRNAs increased significantly (Fig 2C-D). Notably, displacement patterns of differentially expressed piRNAs were dominated by 3’-end shifts, with secondary 5’-end changes (Fig 2E-F). Further analysis of piRNAs with significantly different expression levels compared to the control group revealed more pronounced 3’ end peak shifts, with some shifts at the 5’ end but not as significant as those at the 3’ end (Fig 2G-H). The lengths of downregulated piRNAs were primarily concentrated around 28 nt, while those of upregulated piRNAs were concentrated above 32-36 nt (Fig 2I-J). Strikingly, functional annotations revealed that differentially expressed piRNAs were enriched in exon-derived and rRNA-derived populations, with rRNA-derived piRNAs constituting the largest fraction (Fig 2K-L). A subset of highly upregulated piRNAs originated from aspartic acid tRNAs (Asp-tRNAs), which are distributed across multiple silkworm chromosomes (Fig 2M-N). This aligns with prior evidence that tRNAs contribute to td-piRNA production in silkworms (Honda et al., 2017). Collectively, these findings suggest that SUGP1 regulates piRNA biogenesis by modulating their length, positional bias, and genomic origin. Loss of SUGP1 likely impairs piRNA maturation, leading to truncated or elongated species, which may explain the observed transposon de-repression and viral activation downstream.

**Fig 2.**
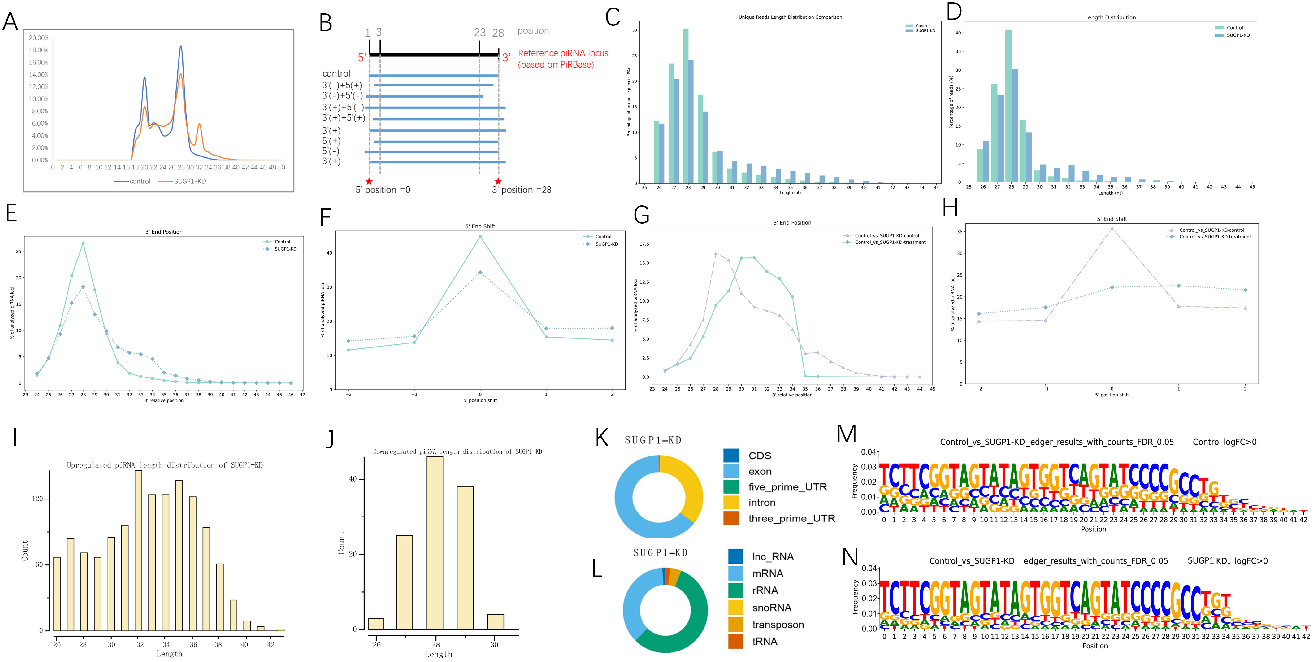
Functional disruption of SUGP1 leads to transposon and vira derepression. (A-B)Changes in BmLV/ BmPnV viral load after knockdown of SUGP1. (C)Amplification of BmLV viral gene fragments in BmE cells of the silkworm. (D-E)Changes in BmLV/ BmPnV viral load after knockdown of core components of the piRNA biosynthetic pathway. (F)Changes in transposon expression after SUGP1 knockdown. (G-I)Changes in Masc/Siwi/BmAgo3 expression after SUGP1 knockdown.

### 0.3 SUGP1 establishes a piRNA biosynthesis pathway through interaction with Y-box protein

SUGP1 dysfunction inhibits of normal piRNA biosynthesis, as evidenced by reduced piRNA abundance and altered length distribution (Fig 2A-N). To further investigate how SUGP1 participates in the piRNA biosynthesis pathway, we constructed a SUGP1 overexpression vector tagged with Flag, using Flag-EGFP as a control (S2 FigA-B). qPCR and WB results confirmed robust SUGP1 expression in BmE cells of the silkworm (S2 FigC-F). Subsequent immunoprecipitation-mass spectrometry (IP-MS) identified proteins interacting with SUGP1. Silver staining revealed that Flag-SUGP1 pulled down distinct protein complexes compared to control Flag-EGFP and IgG (Fig 3A, S1 Table).Mass spectrometry analysis highlighted Y-box protein (Ybp) as the top interactant, a nuclear-cytoplasmic shuttling protein implicated in transcriptional regulation, mRNA splicing, stress granule assembly, and P-body formation (involved in piRNA biosynthesis) [47–50]. Co-immunoprecipitation (Co-IP) and immunofluorescence confirmed physical interactions between SUGP1 and Ybp (Fig 3B,D; S2 FigG; S3 FigA-B). Additionally, interactions were observed between Y-box protein and Nxt1 (a nuclear-cytoplasmic transport factor during piRNA biogenesis) and Spn-E (a nucleolar protein critical for piRNA processing and also present in the P body), consistent with its role in piRNA-related pathways (Fig 3C-D; S2 FigG; S3 FigA-B). To further validate whether the Y-box protein is involved in piRNA biogenesis, we performed RNA-seq and small RNA-seq in Ybp knockdown (Ybp-KD) sample. RNA-seq analysis revealed significant dysregulation of pathways involved in splicing, nuclear-cytoplasmic transport, tRNA biogenesis, and viral infection, aligning with Ybp’s known functions (Fig 4A-B). Consistent with SUGP1-KD effects, Ybp-KD led to reduced 26–29 nt piRNA abundance and increased 31–37 nt piRNA types (Fig 4C-D), with abnormal 3’-end elongation in a subset of piRNAs (Fig 4E-F). These findings suggest that SUGP1 orchestrates piRNA biogenesis via Ybp-dependent recruitment to P-bodies or associated complexes, coordinating nuclear-cytoplasmic trafficking and processing of piRNA precursors.

**Fig 3.**
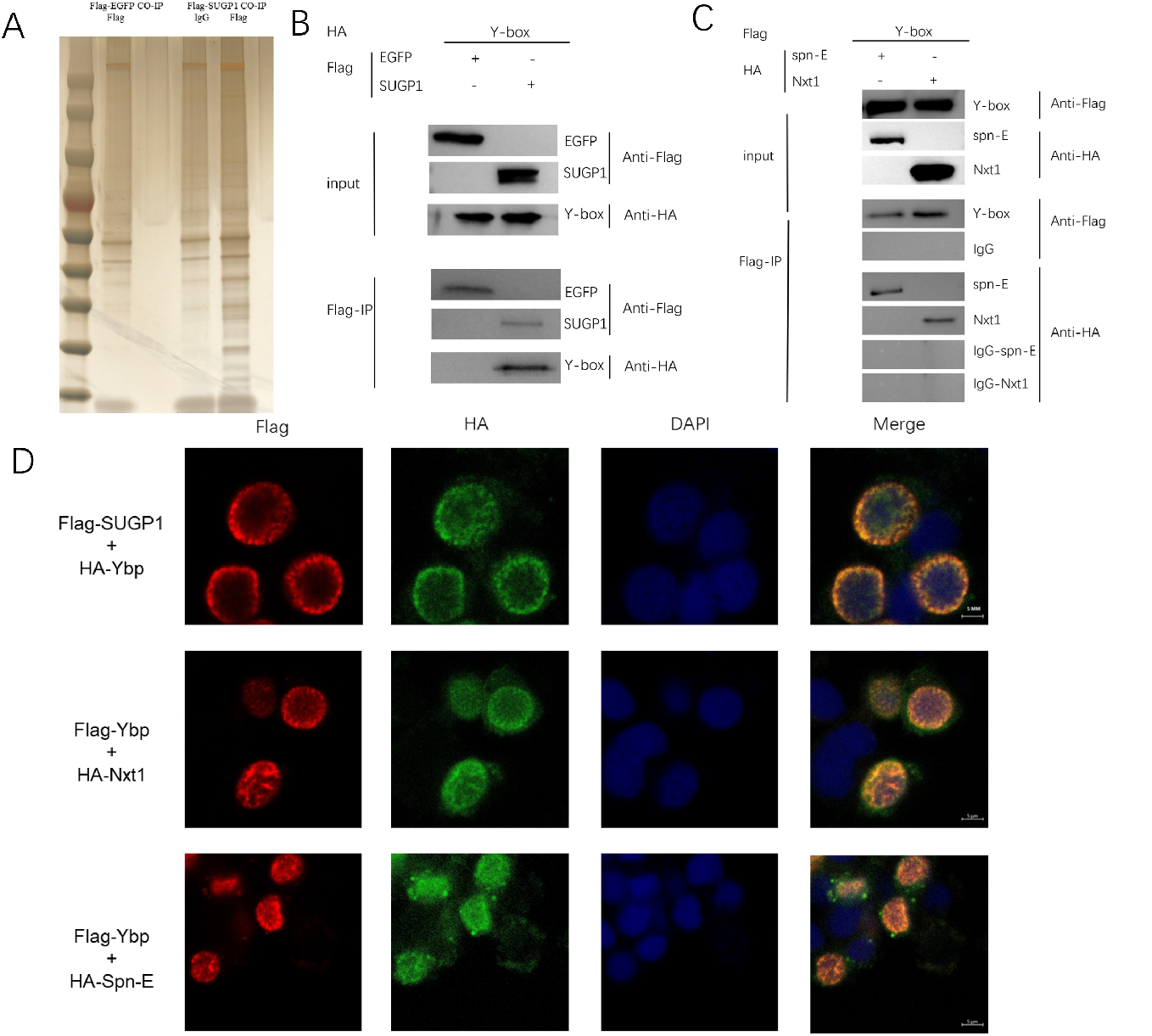
SUGP1 establishes a piRNA biosynthesis pathway through interaction with Y-box protein. (A) Silver staining results of samples after immunoprecipitation. (B) Western blot shows interaction between SUGP1 and Y-box protein. (C) Western blot shows interaction between Y-box protein and Nxt1, Spn-E. (D) Immunofluorescence (200x) showed colocalization between SUGP1 and Y-box protein, Y-box protein and Nxt1, and Y-box protein and Spn-E protein.

**Fig 4.**
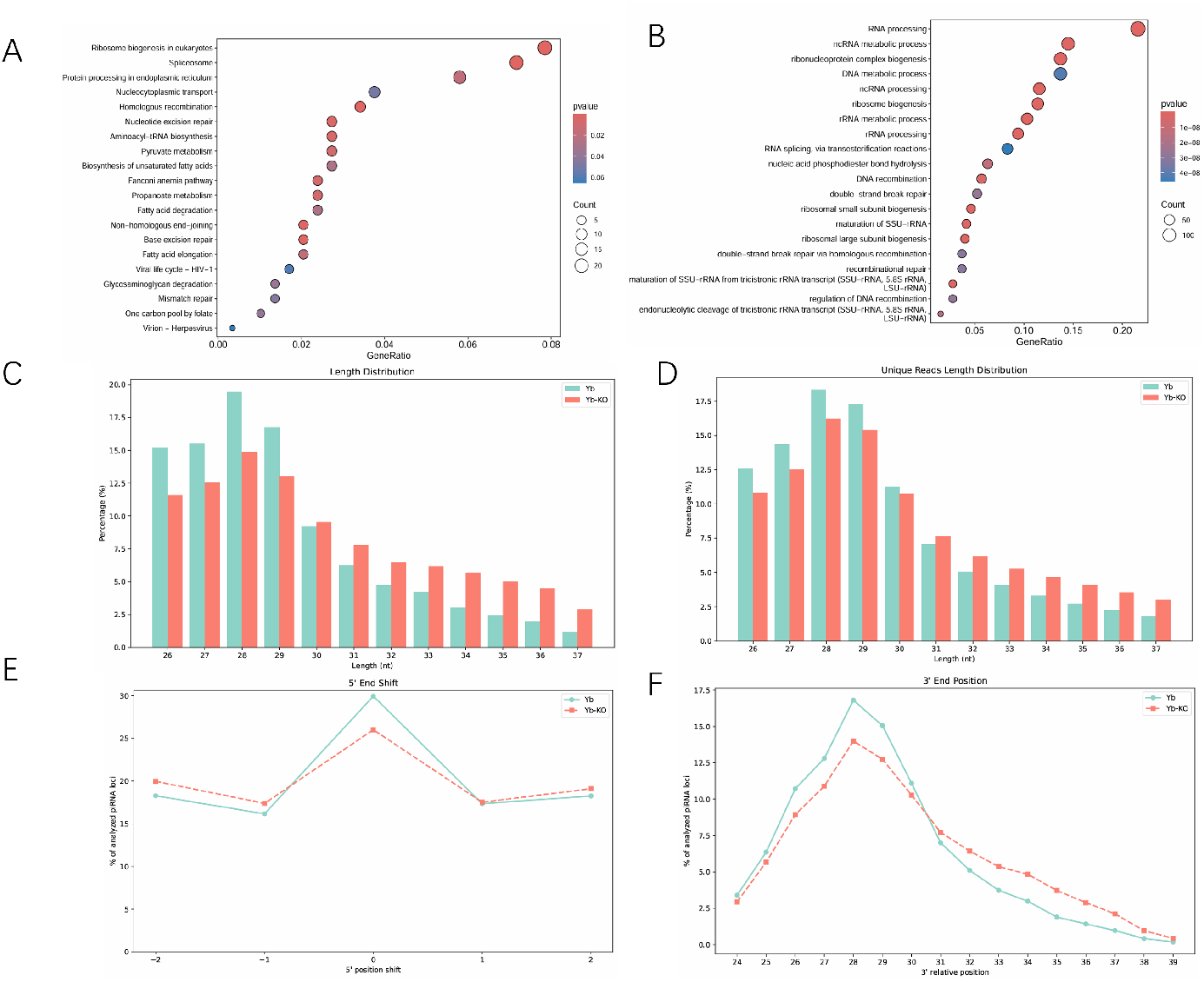
Knockdown of Y-box protein causes abnormalities in piRNA biosynthesis. (A) GO enrichment analysis of differentially expressed genes after Y-box protein knockdown. (B) KEGG enrichment analysis of differentially expressed genes after Y-box protein knockdown. (C-D) The length distribution of piRNAs from 25-45 nt libraries mapped to PiRBase database under knockdown of Y-box protein (Figure C shows the deduplicated piRNA data, representing the types of piRNA; Figure D represents the total amount of all piRNA). (E-F) The 5’-and 3’-end variation analysis of piRNAs from 25-45 nt libraries mapped to PiRBase database under knockdown of Y-box protein. The mode values of 5’- and 3’-ends at each piRNA locus were calculated and the percentage in each length are indicated. Knockdown of Y-box protein extended the 3’ ends of piRNAs but not their 5’ ends.

### 0.4 The role of SUGP1 in piRNA biosynthesis in silkworms is not conserved across species

while our results have demonstrated that SUGP1 modulates piRNA biogenesis in silkworms, its functional conservation across species remains unclear. To verify whether the effect of SUGP1 on piRNA biogenesis is conserved across species, we examined the types, numbers, and displacement changes of piRNAs in SUGP1-RNAi Drosophila melanogaster. In both types of SUGP1-RNAi Drosophila melanogaster, SUGP1 expression levels were reduced by 40-80% (Fig 5A; S4 FigA), concomitantly downregulating piwi expression (Fig 5B). Surprisingly, no significant changes in piRNA abundance or length distribution were observed in SUGP1-RNAi flies, though minor increases in unique piRNA types were detected (Fig 5C-D). 5’ and 3’-end positional shifts of piRNAs remained unchanged (Fig 5E-F), contrasting sharply with silkworm phenotypes. However, SUGP1-RNAi caused striking reproductive defects in D. melanogaster. Egg-to-adult viability dropped significantly, with effects correlating directly with RNAi efficiency (Fig 5G; S4 FigA-D). Notably, pupal morphology (length/weight) remained unaffected (Fig 5H-I; S4 FigC-D), suggesting species-specific functional divergence of SUGP1. While SUGP1’s role in silkworm piRNA biogenesis is central, its conserved function in D. melanogaster may instead prioritize developmental processes unrelated to canonical piRNA pathways.

**Fig 5.**
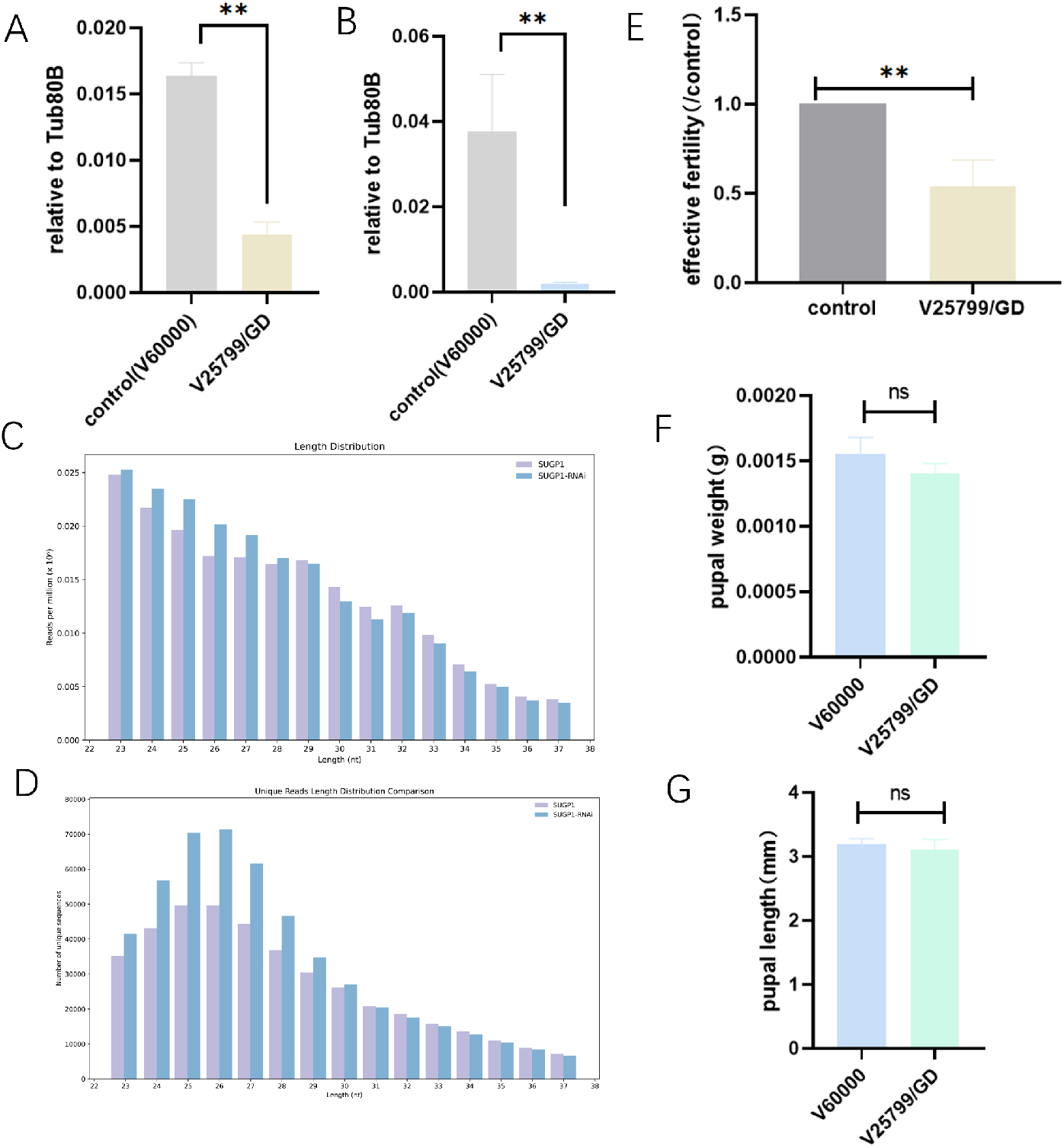
The role of SUGP1 in piRNA biosynthesis in silkworms is not conserved across species. (A) qPCR of SUGP1 after SUGP1-RNAi(V25799/GD). (B) qPCR of piwi after SUGP1-RNAi(V25799/GD). (C-D) The length distribution of piRNAs from 21-39 nt libraries mapped to flyBase database under RNAi of SUGP1(V25799/GD) (Figure C shows the deduplicated piRNA data, representing the types of piRNA; Figure D represents the total amount of all piRNA). (E-G) Changes in pupal length, pupal weight, and effective reproductive capacity in Drosophila following SUGP1-RNAi(V25799/GD).

## Discussion

Genome-wide screening, characterized by its unbiased, high-throughput, and comprehensive nature, represents a paradigm shift from the traditional “hypothesis-verification” model to a “discovery-driven” approach, significantly accelerating our understanding of gene function and disease mechanisms. Prior studies employing RNAi-based genome-wide screens in species such as Drosophila and Caenorhabditis elegans have identified core components of the piRNA biogenesis pathway [34–36]. However, our in-depth analysis of these datasets revealed additional novel candidate genes requiring functional validation, including splicing-related genes that are not conserved across species. This observation prompted us to hypothesize whether splicing factors might also play unappreciated roles in piRNA biogenesis. As a representative model organism for Lepidoptera, the silkworm offers unique experimental advantages in studying the mechanisms of piRNA biosynthesis: 1) silkworm cells possess a complete piRNA biosynthesis pathway, and 2) despite its compact genome, the species exhibits remarkable piRNA diversity [41, 42] . In this study, we focused on SUGP1, a splicing factor critically involved in 3’ splice site recognition and known to interact with SF3B1, a protein frequently mutated in cancer. Recent studies highlight the emerging role of piRNAs in tumorigenesis [38–40], suggesting potential cross-talk between splicing regulation and piRNA function. Furthermore, preliminary experiments revealed that endogenous Masc expression was significantly upregulated upon SUGP1 knockdown, which is consistent with an antagonistic interplay between Masc-mRNA and Fem-piRNA. These observations motivated a multidimensional investigation of whether SUGP1 directly regulates piRNA biogenesis and the underlying mechanisms.

Our RNA-seq analysis of SUGP1-knockdown (KD) cells revealed aberrant splicing events alongside dramatic increases in viral loads (e.g., BmLV and BmPnV), coinciding with accelerated cellular degeneration. Given the well-documented role of piRNAs in antiviral defense [51–53], these findings strongly implicated a functional link between SUGP1 and piRNA pathways. To more directly assess whether SUGP1 is associated with piRNA biosynthesis, small RNA-seq was performed, which revealed SUGP1’s direct influence on piRNA length distribution, abundance, and genomic origin. Notably, this study fills a critical gap in previous screening efforts, which lacked rigorous validation of splicing-related candidates in piRNA biosynthesis.

To further investigate how SUGP1 participates in the piRNA biogenesis pathway, we constructed an Flag-tagged overexpression vector and identified interacting proteins using IP-MS. Notably, the Y-Box protein emerged as the top hit with a remarkably high score, prompting us to explore its functional relevance. A review of relevant literature revealed that Y-box protein have diverse roles in RNA processing and regulatory pathways and are known to share subcellular localization or phase-separated condensates with core piRNA biogenesis machinery [47–50]. This spatial and functional overlap suggests that SUGP1 may bridge splicing regulation and piRNA biogenesis by physically interacting with Y-box proteins. Complementary assays, including Western blotting and immunofluorescence, corroborated these findings, demonstrating co-localization and interaction between SUGP1 and Y-box proteins. Crucially, small RNA-seq analysis following Y-box protein knockdown recapitulated the SUGP1-KD profile, further supporting a conserved mechanistic link between these factors in piRNA production.

In previous studies, the Drosophila splicing regulator RNPS1 was implicated in piRNA biogenesis via Piwi expression modulation [46] . However, our experiments revealed no significant downregulation of silkworm Piwi homologs (Siwi and BmAgo3) in SUGP1-KO cells, suggesting functional divergence between species. To address whether SUGP1’s role in piRNA biogenesis is conserved, we extended our analysis to Drosophila. However, due to technical limitations in rRNA depletion for Drosophila small RNA-seq, we applied stringent quality control and bioinformatic filtering to mitigate background noise. Despite these measures, the small RNA-seq profiles of Drosophila SUGP1-RNAi lines failed to reproduce the silkworm-specific piRNA length/distribution shifts, demonstrating species-specific differences in SUGP1’s functional interaction with the piRNA pathway. This finding underscores the evolutionary plasticity of piRNA biogenesis mechanisms and highlights the silkworm’s unique value as a model for exploring novel regulatory modules distinct from classical Drosophila systems.

In summary, our findings collectively reveal that SUGP1 serves as a pivotal regulator in piRNA biogenesis through its functional interplay with the Y-box protein, bridging nuclear RNA processing and cytoplasmic piRNA maturation. The discovery of this conserved mechanistic link between Y-box proteins and piRNA pathways across species underscores the evolutionary functional convergence of RNA-binding proteins in small RNA systems. However, the striking species-specific divergence observed in SUGP1-dependent piRNA regulation between silkworms and Drosophila highlights the evolutionary plasticity of this machinery. These results not only refine our understanding of piRNA biogenesis complexity but also raise intriguing questions about how developmental and environmental pressures drive adaptive modifications in conserved RNA pathways. Future studies should prioritize dissecting the temporal dynamics of SUGP1/Y-box condensate assembly and exploring comparative genomic approaches to map the genetic determinants underlying species-specific regulatory differences. Ultimately, this work establishes the silkworm as a unique model organism for uncovering non-canonical mechanisms of piRNA biogenesis, offering potential insights into the treatment of diseases linked to piRNA dysregulation, such as transposon hyperactivation in cancer and germ cell disorders.

## Materials and methods

### Construction of plasmid vectors

2× Hieff Canace® PCR Master Mix High Fidelity Enzyme Premix produced by yeasen Biological Company was used to amplify the target fragment by PCR, and Gel Extraction Kit produced by OMEGA was used to recover the target fragment. Restriction endonuclease and T4 DNA ligase produced by NEB Company were used for enzyme digestion and ligase of target fragments, and receptive cell Trans-T1 produced by Transgen company was used for transformation experiment. Endotoxin-free plasmid extraction was performed using QIA prep Spin Mini prep Kit produced by Qiagen.

### Cell culture

The BmE cell line used was obtained from the Biological Science Research Center, Southwest University. Cells were passaged every two days by replacing the medium with Grace insect culture medium supplemented with 10% fetal bovine serum (Gibco, New York, USA), with an average passage interval of 5 days.

### Transfection of BmE cells

The cells in good condition with a density of 80%-90% were spread in the 12-well cell culture plate with a density of about 70%. Cells in 12-well plates were cultured overnight until the density reached 80-90%. Preparation was performed as 1.6 µg plasmid and 4 µL transfection reagent were required per cell well. 200 µL Grace medium (Gibco, New York, USA) was added to a sterile 1.5 mL centrifuge tube and 3 µL Transfection reagents (Roche’s X-treme GENE HP DNA Transfection reagents; Roche, Basel, Switzerland), mixed. 200 µL Grace medium was added to a sterile 1.5 mL centrifuge tube, and 1.6 µg plasmid was added and mixed. The plasmid mixture was added into the transfection reagent mixture and left for 20 min after mixing. Add the total mixture into the culture well and mix well. After 6-8 h at 27°C, the culture medium was completely changed and continued. Follow-up tests were conducted as required.

### The design of sgRNA

The sgRNA required for single gene knockout was designed online using the CCTop website (http://crispr.cos.uni-heidelberg.de). Gene sequence information was obtained from the silkDB3.0 database. NGG was selected for PAM sequence, AAGT was used for the 5’ end joint of justice chain, and AAAC was used for the 5’ end joint of antisense chain. The number of base of seed sequence was 12 bp, and the number of base of maximum mismatch was 4 bp. According to the gRNA score and ranking predicted by the website, oligonucleotide single chains were synthesized from sgRNA with fewer miss sites, high target efficiency and high ranking.

### Real-time fluorescence quantitative PCR (qPCR)

Cell RNA was extracted using HiPure Universal RNA Mini Kit of Magen Bio. RNA reverse transcription was performed using One-Step gDNA Removal and cDNA Synthesis SuperMix of Transgen Bio. The qPCR assay was performed using Top Green qPCR SuperMix (+Dye I) of Transgen Bio.

### Western blot (WB)

Cultured cells were collected and washed twice with PBS. NP40 lysate (Beyotime, Shanghai, China) and Protease inhibitor mixture (Beyotime, Shanghai, China) were mixed at a volume ratio of 100:1, and added to cell samples, then lightly mixed and bathed in ice for 30 min. The sample was centrifuged at 12000 rpm at 4°C for 15 min, and the supernatant was the total protein of the sample. The BCA protein quantitative kit produced by Beyotime Biological Company was used to determine the concentration. The same amount of samples were taken according to the protein concentration, and the appropriate amount of 5×SDS-PAGE Loading Buffer was added. After heating at 95°C for 10 min, SDS-PAGE was performed. After electrophoresis, the protein sample was transferred to PVDF membrane by wet transfer method. 5% skim milk powder was used to seal the membrane. After sealing, 1:5000 times diluted primary antibody (rabbit monoclonal antibody against HA; abclonal, Shanghai, China. mouse monoclonal antibody against Flag; HUABIO, Huangzhou, China. mouse monoclonal antibody against tubulin; Beyotime, Shanghai, China) incubated at room temperature for 1 h and then treated with 1:5000 times diluted secondary antibody (HRP labeled Goat Anti-mouse/ rabbit IgG; Beyotime, Shanghai, China) was incubated at room temperature for 1 h and finally exposed using the Enhanced ECL Chemiluminescent Substrate Kit produced by Yeason Bio.

### Immunofluorescence (IF)

Discard the medium, slowly add room temperature PBS on the cells, wash 3 times, 5min each time; on the cells covered with 4% neutral formaldehyde fixative, fixed at room temperature for 15min; pre-cooled PBS buffer at 4 °C, wash 3 times, 5min each time; 0.3% Triton™ X-100 10min; 5% goat serum(Beyotime, Shanghai, China) will be completely covered with samples at 37 °C constant temperature and humidity incubator incubation 30min; remove the sealing solution, directly on the sample dropwise addition of PBS buffer prepared primary antibody (rabbit monoclonal antibody against HA; abclonal, Massachusetts, USA. mouse monoclonal antibody against Flag; HUABIO, Huangzhou, China) working solution, placed in 4 °C shaking bed incubation overnight, the next day the sample at room temperature, rewarming 15min after removing the antibody working solution, washed with buffer PBS 3 times, 5 minutes each time; in the sample dropwise addition of fluorescent secondary antibody (Cy3 labeled Goat Anti-mouse IgG(H+L), FITC labeled Goat Anti-rabbit IgG(H+L); Beyotime, Shanghai, China) working solution corresponds to the species of the primary antibody, the sample needs to be completely covered, protected from light, 37 °C, shaking bed incubation for 1 hour; remove the secondary antibody working solution, wash with buffer 1x TBS 2 times, 5 minutes each time; add 1µg/mL DAPI (Beyotime, Shanghai, China) working solution dropwise to the sample, avoid light, and incubate at room temperature on a shaking bed for 10 minutes; remove the DAPI working solution, and wash with buffer PBS 3 times, 5 minutes each time; remove the cell crawler, and then cover it on the slide titrated with the anti-fluorescence attenuating sealer and observe and collect the image under a fluorescence microscope. After the cell crawls were removed and covered with a drop of anti-fluorescence attenuation sealer, they were observed under a confocal microscope and images were collected.

### Co-immunoprecipitation (CO-IP)

Take 40ul of DynabeadsTM protein G well-mixed magnetic beads (ThermoFisher, Waltham, USA) and wash the magnetic beads with 100ul of NP40(Beyotime, Shanghai, China) lysate containing protease inhibitor (100:1) on a rotator 3 times (1min/time); 100ul of NP40+antibody containing protease inhibitor (control: Ig G; antibody to the target: 2-4ul, usually 3ul), incubate for 30min-40mim on a rotator, and then wash the magnetic beads 3 times with NP40 with protease inhibitor; add 200ul of crosslinking agent (1ml of antigntion buffer+0.0029g BS3), incubate on a rotator for 30min-40mim; add 12.5ul of Quending buffer (1M Tris-HCl), incubate for 15min; add NP40 to wash the magnetic beads for 3 times; then add protein samples, rotate and incubate at 4°C for 6h; discard the supernatant, wash the magnetic beads for 3 times with PBS; add 100-120ul of SDT eluent (SDT :4% SDS in 50mM tris-HCl), incubate for 5min.

### Silver dye

After electrophoresis, take the gel and put it into 100ml fixed solution (50ml ethanol, 10ml acetic acid and 40ml Milli-Q pure water), and shake it for 40 minutes at room temperature on a 60-70rpm shaker; After fixation, shake with 30% ethanol at room temperature for 10 minutes on a shaker at 60-70rpm; Shake 200ml of Milli-Q grade pure water at room temperature on a shaker at 60-70rpm for 10 minutes; After complete washing, discard the water and add 100ml of silver dye sensitization solution (1X). Shake at room temperature for 2 minutes on a shaker at 60-70rpm; Discard the original solution, add 200ml of Milli-Q grade pure water, and shake at room temperature for 1 minute (2 times) on a shaker at 60-70rpm. Then discard the water, add 100ml of silver solution (1X), and shake at room temperature for 10 minutes on a shaker at 60-70rpm; Afterwards, discard the original solution and add 100ml of Milli-Q grade pure water. Shake on a shaker at room temperature for 1.5 minutes; After the silver staining is completed, add 100ml of silver staining solution and shake at room temperature for 3-10 minutes on a shaker until an ideal protein band appears. Then discard the silver staining solution and add 100ml of silver staining termination solution (1X). Shake at room temperature on the shaker for 10 minutes; After achieving significant color development, discard the silver dye termination solution and add 100ml of Milli-Q grade pure water. Shake on a shaker at room temperature for 5 minutes. All silver staining premix reagents are sourced from Biyuntian P0017S (Beyotime, Shanghai, China).

### Graphing Software

Fluorescence intensity, length, and other quantitative measurements in images were processed using ImageJ. Quantitative data visualization was achieved with GraphPad Prism. GO and KEGG diagrams utilized the online platform CNS(https://cnsknowall.com/index.html/HomePage)

### Small RNA-seq

The total RNA of the samples was extracted using the MJZol total RNA extraction kit (Majorbio, China). For the enriched small RNA fragments of 16 - 35 nt, the QIAseq miRNA Library Kit (Qiagen; Germany) was used to construct the library, and sequencing was performed using the SE75 sequencing mode on the illumina navoseq Xplus platform.

### RNA-seq

Extract the total RNA of the sample ((USA, life, No.: 10296010)) and digest the DNA with DNaseI. Then, use the Agilent 2100 Bioanalyzer (USA, Agilent Technologies, Inc.) and the ultraviolet spectrophotometer NanoDrop (USA, Thermo scientific) to detect the purity of the RNA, and test the concentration, purity, etc. of the RNA sample. After the test results meet the requirements, use Oligo(dT) magnetic beads + hexamer random primers for library construction. Use the Agilent 2100 Bioanalyzer to detect the insert fragment range of the library and use the ABI StepOnePlus RealTime PCR System (USA, Applied Biosystems, Inc., No.: 4376600) to quantify the concentration of the library. After the quality inspection is qualified, use the MGI T7 (China, MGI, Inc) sequencer for sequencing.

### Small RNA Sequencing Analysis

#### 0.4.1 Data Preprocessing and Quality Control

Raw small RNA sequencing reads were subjected to a rigorous quality control workflow. Initial quality assessment was performed using FastQC (v0.11.9). Subsequently, fastp (v0.24.0) was employed to trim the 3’ adapter sequence (5’-AACTGTAGGCACCATCAAT-3’) and filter the reads. Reads were discarded if they had a Phred quality score below 20, contained any ambiguous ‘N’ bases, or fell outside the 18–37 nucleotide length range. The quality of the final clean dataset was confirmed using FastQC and MultiQC (v1.14).

#### 0.4.2 piRNA Identification and Characterization

To identify piRNAs and characterize their properties, clean reads were first converted from FASTQ to FASTA format using seqkit (v2.10.0). Identical reads were then collapsed into unique sequences with their corresponding read counts using fastx collapser (FASTX-Toolkit v0.0.14). These unique sequences were aligned to the Bombyx mori reference piRNA database (piRBase) using seqmap (v1.0.13) under highly stringent conditions, allowing only perfect, forward-strand matches (zero mismatches).Based on these alignments, we first performed a global characterization of the piRNA populations. The overall length distribution for each sample was plotted to assess general size profiles. Furthermore, to investigate potential post-transcriptional modifications, we analyzed nucleotide variations at the 5’ and 3’ termini of the mapped piRNAs relative to their reference sequences. The distribution of these terminal variations was compared between treatment and control groups to identify any systemic shifts in piRNA processing. The alignments also generated a count matrix of all confirmed piRNAs, which served as the basis for subsequent differential expression analysis.

#### 0.4.3 Differential Expression Analysis of piRNAs

Differential expression analysis between experimental and control groups was performed using a custom R script leveraging the edgeR package (v3.34.1). A pairwise comparison approach was employed for each treatment versus control. piRNAs were considered significantly differentially expressed if they met the criteria of a False Discovery Rate (FDR) less than 0.05 and a total read count of at least 100 across the compared samples. The length distributions of both upregulated and downregulated piRNAs were plotted to observe any size preferences.

#### 0.4.4 Characterization of Differentially Expressed piRNAs

To further characterize the differentially expressed piRNAs (DE-piRNAs), we analyzed their sequence motifs and genomic origins. Sequence logos for upregulated and downregulated piRNA populations were generated using a custom Python script to identify conserved nucleotide patterns. For genomic origin analysis, DE-piRNA sequences were aligned to the silkworm reference genome (NCBI: ASM3026992v2) using Bowtie. The resulting alignments were converted to BED format, and intersectBed (bedtools v2.30.0) was used to determine the overlap between DE-piRNA loci and annotated genomic features (e.g., CDS, UTR, intron, exon). The distribution of these features was then quantified and plotted to reveal the primary sources of the DE-piRNAs.

### Transposon Expression Analysis

#### 0.4.5 Transposon Library Construction

To facilitate transposon analysis, an up-to-date transposable element (TE) library was constructed. TEs were identified de novo from the latest silkworm reference genome assembly (NCBI: ASM3026992v2) using HiTE (v3.3.3).

#### 0.4.6 Quantification of Transposon Expression

Using this custom TE library, the expression levels of transposons were quantified from a separate paired-end RNA-seq dataset. Raw reads were processed for quality control using fastp, where adapters were trimmed and low-quality reads were filtered based on multiple criteria, including mean quality score, ‘N’ base content, and complexity. Clean reads shorter than 50 bp were discarded. The expression analysis was then performed by aligning the clean reads to the reference genome using STAR (v2.7.11b) and quantifying TE expression with the TEcount algorithm from the TEtranscripts software package (v2.2.3).

## 1 code availability

All the data analysis code used in the article is available at the code repository ()

## 2 Data availability

The whole genome resequencing data used in this paper are publicly available under accession number PRJNA1337577 in NCBI.

## 3 CRediT authorship contribution statement

LJ and SM conceived the study and designed the experiments. DL constructed the reporter cell line and performed the library construction and screening with the help from RLW and PW. YL designed the reporter construct. YKG performed the bioinformatic analysis. LS provide some advises during the experimental design. DL wrote the manuscript. All authors read and approved the final manuscript.

## 4 Acknowledgments

This work was supported in part by the National Key Research and Development Program of China (grant number 2023YFD1700700), the National Natural Science Foundation of China (grant number 32570591), and the Chongqing Natural Science Foundation (grant number CSTB2024NSCQ-JQX0018).

## Supporting information

**S1 Fig. Abnormal SUGP1 expression leads to transposon and viral de-repression**.(A-E) Several major alternative splicing changes after SUGP1 knockdown. (F) Abnormal splicing products of Bub1b after SUGP1 knockdown. (G) Cell status after SUGP1 knockdown: obvious vacuolation and sparse appearance. The left side is the empty vector transfection control, and the right side is the SUGP1-KD experimental group. (H) Sequence peak diagram of target sites after SUGP1 knockdown: Overlapping peaks appear after the sgRNA site. (I-K) Changes in BmLV/ BmPnV viral expression following SUGP1 knockout as determined by RNA-Seq sequencing.

**S2 Fig. Construction of SUGP1 overexpression vector**. (A) Schematic diagram of tagged overexpression vector construction. (B) Agarose gel electrophoresis of the products obtained after restriction enzyme digestion of the vector backbone. The control is the original plasmid, and the required backbone after digestion is marked with a red frame. (C-D) qRT-PCR results after transfection of cells with overexpression vectors. (E-F) Western blot results after transfection of cells with overexpression vectors (E: Flag-EGFP; F: Flag-SUGP1). (G) Fragment amplification results of related vector construction.

**S3 Fig. SUGP1 establishes a piRNA biosynthesis pathway through coupling with Y-box protein**. (A) Immunofluorescence (40x) showed colocalization between SUGP1 and Y-box protein, Y-box protein and Nxt1, and Y-box protein and Spn-E protein. (B) The fluorescence intensity of fluorescence colocalization in Figure 3D was quantified using ImageJ.

**S4 Fig. The role of SUGP1 in piRNA biosynthesis in silkworms is not conserved across species**. (A) qPCR of SUGP1 after SUGP1-RNAi(V110279/KK). (B) q-PCR of piwi after SUGP1-RNAi(V110279/KK). (C-F) Changes in pupal length, pupal weight, and effective reproductive capacity in Drosophila following SUGP1-RNAi(V110279/KK). The interference efficiency of the V110279/KK strain is lower than that of V25799/GD, so its effect on piwi and effective reproductive capacity is weaker than that of V25799/GD.

**S1 Table. IP-MS results**.

